# Systems-level analyses dissociate genetic regulators of reactive oxygen species and energy production

**DOI:** 10.1101/2023.10.14.562276

**Authors:** Neal K. Bennett, Megan Lee, Adam L. Orr, Ken Nakamura

**Affiliations:** Gladstone Institute of Neurological Disease, Gladstone Institutes, San Francisco, CA, 94158, USA; Aligning Science Across Parkinson’s (ASAP) Collaborative Research Network, Chevy Chase, MD, 20815; Appel Alzheimer’s Disease Research Institute, Weill Cornell Medicine, New York, NY, USA; Feil Family Brain and Mind Research Institute, Weill Cornell Medicine, New York, NY, USA; Graduate Programs in Neuroscience and Biomedical Sciences, University of California San Francisco, San Francisco, California, USA; Department of Neurology, University of California, San Francisco, San Francisco, California, 94158, USA

## Abstract

Respiratory chain dysfunction can decrease ATP and increase reactive oxygen species (ROS) levels. Despite the importance of these metabolic parameters to a wide range of cellular functions and disease, we lack an integrated understanding of how they are differentially regulated. To address this question, we adapted a CRISPRi- and FACS-based platform to compare the effects of respiratory gene knockdown on ROS to their effects on ATP. Focusing on genes whose knockdown is known to decrease mitochondria-derived ATP, we showed that knockdown of genes in specific respiratory chain complexes (I, III and CoQ10 biosynthesis) increased ROS, whereas knockdown of other low ATP hits either had no impact (mitochondrial ribosomal proteins) or actually decreased ROS (complex IV). Moreover, although shifting metabolic conditions profoundly altered mitochondria-derived ATP levels, it had little impact on mitochondrial or cytosolic ROS. In addition, knockdown of a subset of complex I subunits—including NDUFA8, NDUFB4, and NDUFS8—decreased complex I activity, mitochondria-derived ATP and supercomplex level, but knockdown of these genes had differential effects on ROS. Conversely, we found an essential role for ether lipids in the dynamic regulation of mitochondrial ROS levels independent of ATP. Thus, our results identify specific metabolic regulators of cellular ATP and ROS balance that may help dissect the roles of these processes in disease and identify therapeutic strategies to independently target energy failure and oxidative stress.

**Significance:** Mitochondrial respiration generates both energy (ATP) and reactive oxygen species (ROS). Insufficient energy and increased ROS from respiratory chain dysfunction may be central to the pathophysiology of neurodegenerative diseases and aging. We established a screening platform using CRISPR and fluorescent-cell sorting to compare the impact of decreasing respiratory chain proteins on ROS and ATP levels. The results provide the first systems-level analysis of how ROS and ATP are differentially regulated, and identify genes and respiratory chain complexes that can manipulate each independently. These findings advance our understanding of the relative contributions of ATP and ROS to disease pathophysiology, and guide the development of therapies to preserve energy while minimizing ROS.

## Introduction

The mitochondrial respiratory chain plays a central role in energy production, and respiratory chain dysfunction leading to energy failure has been implicated in the pathophysiology of numerous inherited and age-associated diseases, including neurodegenerative conditions like Parkinson’s disease and Alzheimer’s disease. However, the relative impact of respiratory chain dysfunction on energy failure versus its other critical functions, including the generation of reactive oxygen species (ROS), regeneration of matrix NAD+, and metabolite biosynthesis, remains less well understood. In particular, excessive ROS production from respiratory chain dysfunction has been widely proposed to cause many of the same diseases in which energy failure is implicated. However, it is challenging to dissociate the impact of mitochondrial effects on energy from those on ROS. As such, it remains unclear which respiratory chain functions are most critical for physiologic function, and which ones, when compromised, drive disease.

Mitochondrial ATP production is driven by the proton-motive force across the inner mitochondrial membrane, which is dependent on electron transport through the respiratory chain. Multiple sites in the respiratory chain, including in complex I and complex III, directly reduce oxygen to generate ROS, primarily superoxide and hydrogen peroxide(1). Cellular ROS levels are highly regulated to facilitate ROS-mediated cellular signaling(2, 3) while mitigating ROS-induced damage to proteins, lipids, and DNA. Although ATP and ROS production are both produced in the course of respiration, the degree to which they are interdependent is poorly understood.

The enzymatic activity of respiratory complexes is regulated by a number of factors, including complex I conformation(4, 5), and complex subunit transcription(6-8) or degradation(9). However, the functional effects of loss of individual subunits is not well understood. Respiratory chain efficiency is also believed to be regulated through the formation of a range of stoichiometric combinations of complexes, termed supercomplexes, with evidence suggesting that supercomplexes promote more ATP and less ROS production than independent respiratory complexes(10-12). However, it is unclear whether all supercomplexes display this bioenergetic efficiency, and the extent to which the expression level of individual subunits can influence supercomplex formation is also unknown.

Although select pharmacologic tools can help dissociate the relative impact of respiratory chain function on ROS and ATP production, systems-level genetic analyses have not been performed. To begin to address this, we previously developed a CRISPR- and FACS-based screening platform to identify genetic modulators of ATP, and used it to define the “ATPome”, a compendium of genes and pathways that regulate ATP levels(13). As expected, this analysis revealed numerous respiratory chain genes that are required to maintain mitochondrial-derived ATP levels but, on its own, did not provide insight into the impact of these same genes on ROS. Here, we adapt our screening approach to detect ROS, and use it to identify genes and pathways that differentially impact ATP and ROS. This work identifies genetic disease targets in which energy and oxidative stress become decoupled, and whose severity may be modulated by either antioxidant or energetic therapy.

## Results

### A robust, antioxidant-sensitive screen for genetic regulators of ROS

We previously developed a screening approach to identify genetic regulators of ATP level, defining the “ATPome”. From this we generated a mini-library of 431 sgRNAs targeting 200 ATP-regulating genes, including 91 primarily mitochondrial genes that decreased ATP in respiratory conditions, and 19 non-targeting sgRNAs as controls. To investigate if our screening approach could be used to identify genetic regulators of ROS, we pre-incubated K562 cells expressing dCas9-KRAB and this CRISPRi sgRNA mini-library for 30 minutes with a ROS-sensitive dye, either MitoSOX Red to detect primarily mitochondrial superoxide(14), DCFDA for general cytosolic ROS including peroxides(15, 16) (Figure 1A), or MitoNeoD, a mitochondria-targeted superoxide probe (17). These incubations were performed in the presence of respiratory metabolic substrate (10 mM pyruvate and 10mM 2-deoxyglucose) to force the cells to rely on mitochondria for energy. Cells were then sorted based on ROS fluorescence signal to isolate cells from the highest or lowest quartiles. Sorted fractions were sequenced to quantify relative representation of each CRISPRi sgRNA in the high-versus low-ROS cell populations, and to calculate a ROS phenotype. Phenotype z-scores were calculated relative to the distribution of pseudogenes, calculated from non-targeting sgRNA(13). This approach robustly determined ROS phenotypes associated with individual gene knockdowns (Figure 1B, S1A, Table S1, Dataset S1).

**Fig. 1.**
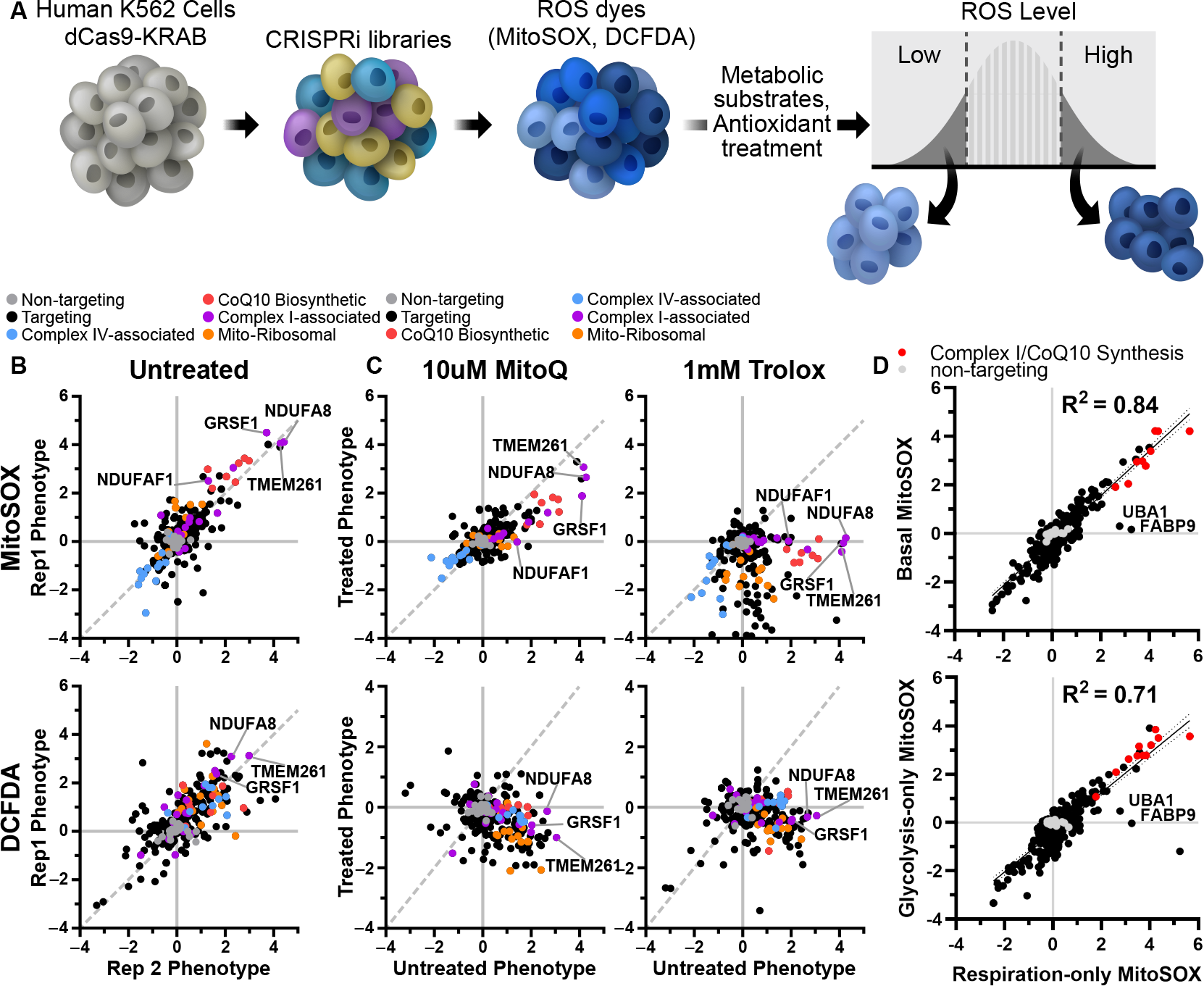
CRISPR screen for genetic regulators of ROS finds robust, antioxidant-sensitive ROS phenotypes. A) Schematic describing a mini-library of CRISPRi sgRNAs that identifies ROS phenotypes based on MitoSOX or DCFDA levels via FACS. B) K562 cells expressing a CRISPRi mini-library and incubated in respiratory conditions for 1 h prior to cell sorting on MitoSOX or DCFDA staining to measure mitochondrial or cytosolic ROS phenotypes, respectively. Several knockdowns associated with complex I (TMEM261, NDUFA8, GRSF1, NDUFAF1) and CoQ10 biosynthesis (PDSS1, PDSS2, COQ5, COQ2) robustly increase mitochondrial or cytosolic ROS. Complex IV-associated gene knockdowns (COX18, COX16, COX11) robustly decrease mitochondrial ROS. C) High mitochondrial and cytosolic ROS phenotypes associated with complex I knockdown are lowered with antioxidant treatments, either 10 µM MitoQ or 1 mM Trolox. Data compiled from n = 2 experiments. D) ROS phenotypes exhibited high correlation between metabolic substrate conditions. Data compiled from n = 2 replicates for basal, 3 replicates for respiration-only and glycolytic-only.

Overall, most high and low ATP hits did not impact either MitoSOX or DCFDA levels. However, knockdown of a subset of low ATP hits, especially those targeting mitochondrial complex I, increased ROS as measured by all three ROS probes. Among the mitochondrial hits, knockdown of complex I subunit NDUFA8, complex I assembly factor TMEM261(18), and complex I function regulator GRSF1(19) reproducibly induced the highest mitochondrial superoxide and cytosolic ROS levels. Knockdown of CoQ10 biosynthesis enzymes (e.g. COQ2, COQ5, PDSS1, PDSS2) was also associated with increased mitochondrial ROS levels. In contrast, knockdown of complex IV-associated genes (e.g. COX16, COX11, and COX18) actually decreased mitochondrial ROS levels. Most low ATP hits targeting mitochondrial ribosomal genes did not increase either mitochondrial or cytosolic ROS. Low ATP hits that increased cytosolic ROS without affecting mitochondrial ROS included PTCD1, a mitochondrial ribosomal assembly factor and AD-associated risk gene(20), and PTCD3, a related mitochondrial ribosomal protein in which mutations can cause Leigh syndrome(21). Overall, this analysis reveals individual genes with particularly large contribution to mitochondrial ROS pools, including disease-related genes whose dysfunction may be closely entwined with changes in cellular energy and specific subcellular pools of ROS.

Although widely used as relatively specific probes for ROS, both MitoSOX and DCFDA fluorescence can also be altered independently of ROS to generate non-specific oxidation products, and can have non-linear signal with ROS due to oxygen or pH sensitivity (22, 23). Therefore, to validate the specificity of ROS-modifying effects, we assessed ROS signals following co-incubation with antioxidants. Indeed, treatment with the water-soluble vitamin E analog Trolox(24), and to a lesser extent the mitochondrially targeted antioxidant MitoQ(25), markedly decreased the effects of high ROS hits on MitoSOX and DCFDA fluorescence relative to treated controls (Figure 1C, Figure S1B-F, Table S1, Dataset S1). In particular, Trolox showed broad, potent antioxidant efficacy, significantly decreasing mitochondrial superoxide and cytosolic ROS independent of targeting sgRNA and particularly in cells with knockdown in CoQ10 biosynthetic genes (Figure S1F). These observations are consistent with the known antioxidant activities of MitoQ and Trolox: MitoQ abolishes mitochondrial superoxide-induced signaling, whereas Trolox exhibits broader potency(26).

As expected, MitoQ decreased ROS in cells with impaired CoQ10 synthesis. Surprisingly, however MitoQ treatment of cells with knockdown of HK2 or VDAC1 induced increased levels of ROS. Higher concentration of MitoQ can inhibit respiration (27), therefore, we also screened for genes that regulate mitochondrial ROS, measured by MitoSOX, in cells treated with 0.1 µM MitoQ, and compared with cells treated with 0.1 µM decyl-TPP, which contains only the mitochondrial-targeting moiety and not the ubiquinone moiety (Figure S1B-D). We observed that this lower dose of MitoQ was even more selective to lowering ROS levels in cells with impaired CoQ10 synthesis. We hypothesize that the increased levels of ROS seen with higher MitoQ concentration is due to the HK2 knockdown-induced increase in TCA activity (13). The combined effects of increased electron flow and decreased respiration in the setting of higher MitoQ may lead to increased ROS. Furthermore, NDUFAF1 knockdown was rescued at both MitoQ concentrations, while high ROS seen with NDUFA8 knockdown was only rescued at the higher MitoQ concentration, showing that the dose-requirement depends on the specific complex I deficiency. This data also raises the possibility that complex I deficiencies driven by NDUFAF1 could be treated by MitoQ, in order to decrease ROS, although further investigation is required.

Previously, we found that the effect of many gene pathways on ATP levels was dependent on the metabolic substrates available to the cells (Figure S1G)(13). To determine if metabolic substrate had a similar effect on ROS phenotype, we repeated the ROS screen in cells incubated in the same three distinct conditions(13, 28): respiration-only (10 mM pyruvate and 10 mM 2-deoxyglucose), glycolysis-only (2 mM glucose, 3 mM 2-deoxyglucose, and 5 µM oligomycin) or respiration and glycolysis (10 mM glucose and 5 mM pyruvate). Surprisingly, the impact of gene knockdown on mitochondrial superoxide phenotypes was largely independent of changes to cellular metabolism, and the degree of ROS changes among these three substrates was highly correlated (Figure 1D, S1G). These findings show that, in many cases, ATP production can be dissociated from ROS production depending on the metabolic context, and understanding such differences may ultimately help determine the relative contributions of insufficient ATP and excessive ROS production in disease.

Notable exceptions to these substrate-independent correlations in ROS phenotypes were ubiquitin like modifier activating enzyme 1 (UBA1) and fatty acid binding protein 9 (FABP9), which increased mitochondrial superoxide when cells were forced to rely on respiratory metabolic substrates but not in the other two conditions where glycolysis was active (Figure 1D). We speculate that knockdown of these proteins may alter protein and lipid homeostasis, which could contribute to mitochondrial dysfunction and increased ROS. FABP9 is believed to have binding affinity with long-chain fatty acids(29), and deficiencies in mitochondrial metabolism of long-chain fatty acids can cause elevated ROS and severe cardiomyopathy(30). Importantly, these results measuring mitochondrial ROS following UBA1 knockdown indicate that increases in mitochondrial ROS that come with ubiquitinylation defects can be lowered by shifting away from respiratory metabolic substrates.

### Distinguishing ATP and ROS phenotypes

To comprehensively profile the impact of gene knockdown on ROS levels in a broader set of genes, we performed our ROS screen on an expanded library of CRISPRi sgRNAs comprising 54,959 sgRNAs targeting 4,565 genes enriched for mitochondrial genes, encompassing most of the subunits of the electron transport chain, along with 2,800 non-targeting sgRNA controls. We examined ROS levels specifically in respiratory conditions, where we had the most robust ATP phenotypes. As expected, many of the strongest high ROS phenotypes from the smaller library were replicated in the expanded library, for example, NDUFA8 and GRSF1 (MitoSOX phenotype z-scores: 7.40 and 8.22, respectively) (Figure 2A, Dataset S2).

**Fig. 2.**
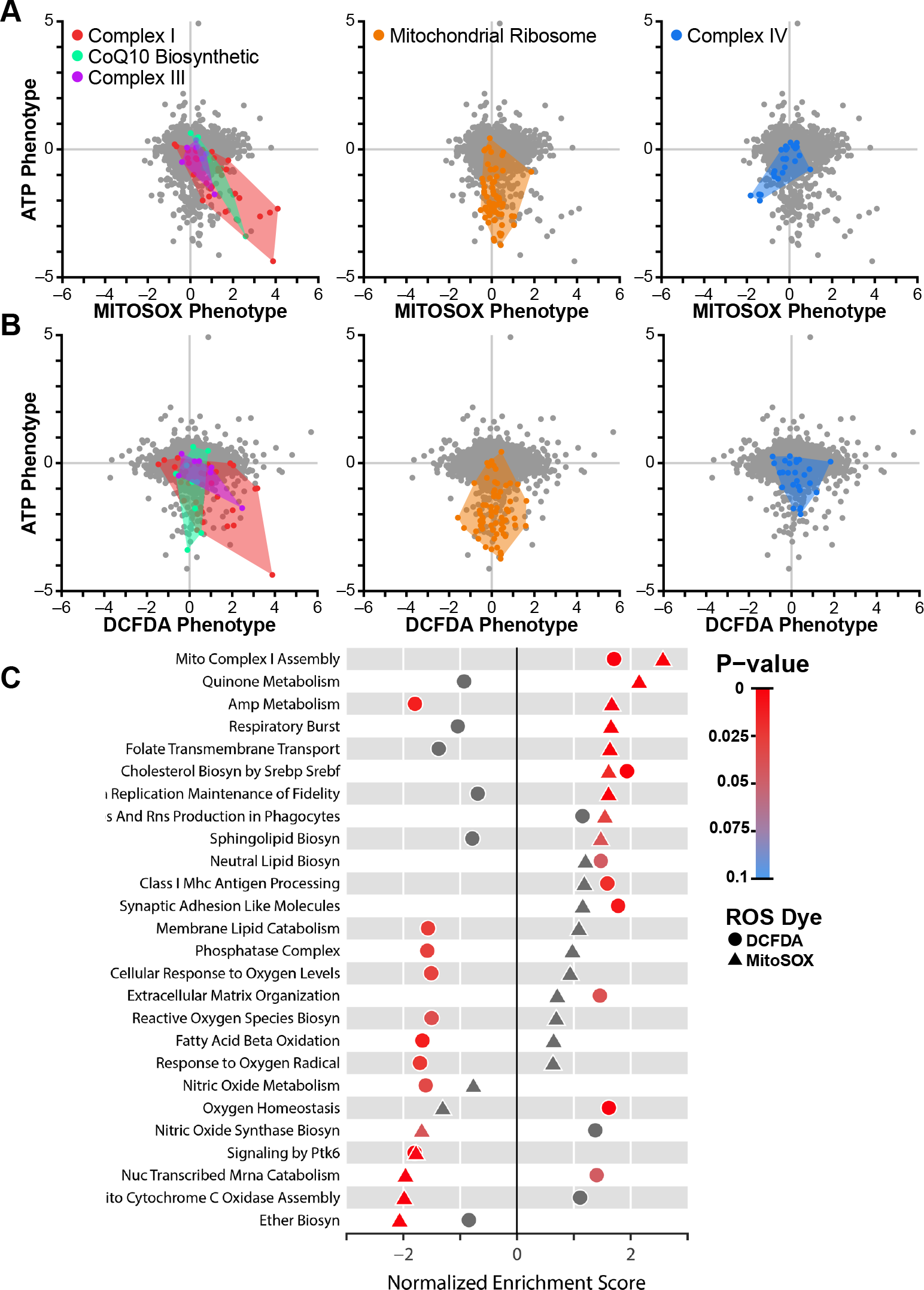
Systems-level analysis of ATP and ROS phenotypes. A) Mitochondrial ROS level assessed in K562 cells expressing an expanded library of CRISPRi sgRNAs with MitoSox or (B) cytosolic ROS with DCFDA were plotted against previously determined ATP phenotypes (Bennett et al.(13)). Knockdown of complex I subunits in the respiratory chain decreased ATP and increased mitochondrial and cytosolic ROS. Knockdown of CoQ10 biosynthetic genes and complex III genes also increased mitochondrial ROS. In contrast, knockdown of mitochondrial ribosomal proteins (orange) strongly decreased ATP, but did not impact ROS, while knockdown of complex IV-associated genes (blue) decreased ATP and mitochondrial ROS. C) Fast, pre-ranked geneset enrichment analysis reveals pathways of genes that regulate either mitochondrial or cytosolic ROS. Besides the respiratory chain-associated pathways, this analysis highlights ether lipid biosynthesis, which occurs in the peroxisome, as a significant regulator of mitochondrial ROS. Data compiled from n = 2 experiments.

Analysis of the expanded library showed that only a subset of hits that increased mitochondrial ROS also increased cytosolic levels. This included certain complex I subunits (NDUFA8, NDUFC1, NDUFAB1, DCFDA phenotype z-scores: 5.36, 4.24, 4.38, respectively) and one complex III subunit (UQCRB, DCFDA phenotype z-score: 3.39) (Figure 2B, Dataset S2).

One interpretation of these results is that individual complex I and III subunits differ in their contributions to mitochondrial versus cytosolic ROS levels. On a pathway level, knockdown of complex I-associated subunits as well as genes involved in cholesterol biosynthesis increased both mitochondrial superoxide and cytosolic ROS, while gene knockdowns in other pathways had divergent ROS phenotypes, particularly those related to AMP metabolism (e.g. APRT, AK4, ADSL, AK2), where mitochondrial ROS levels were increased while cytosolic levels were decreased (Figure 2C). Defects in these AMP metabolism genes are known to cause neurodevelopmental disorders(31). Notably, knockdown of peroxisomal ether lipid biosynthesis genes specifically resulted in low mitochondrial superoxide levels.

Knockdown of nearly all nuclear encoded respiratory chain subunits revealed that many respiratory chain deficits decreased ATP levels, but only a subset of these increased mitochondrial superoxide levels (Dataset S2). Consistent with the mini-library results (Figure 1B), knockdown of mitochondrial ribosomal genes, which results in low ATP phenotypes, did not alter mitochondrial superoxide (Figure 2A, Dataset S2). Similarly, while knockdown of a subset of subunits of complex I, complex III, and CoQ10 biosynthetic genes both decreased ATP and increased mitochondrial superoxide, knockdown of complex IV assembly factors COX11, SCO1, COX16 and COX18 resulted in low mitochondrial superoxide despite also decreasing ATP (MitoSOX phenotype z-scores: -6.55, -4.82, -5.12, -4.98, respectively) (Figure 2A, Dataset S2). Since complex I and complex III contain known sites of mitochondrial ROS generation(1), knockdown of subunits within these complexes may disrupt electron transport, causing ROS production to increase. In contrast, complex IV destabilization can destabilize complex I(13, 28), potentially leading to decreased ROS production upstream of complex IV.

### Identification of critical subunits for complex I bioenergetic function

Complex I deficiency has been widely implicated in the pathogenesis of mitochondrial diseases, Parkinsonism, diabetes, and aging(32-34). However, the relative contribution of increased ROS or decreased ATP production to pathogenesis may vary between diseases. We observed that knockdown of individual complex I subunits resulted in a range of ATP and ROS phenotypes (Figure 3A). Knockdown of NDUFA8 resulted in the largest increase in mitochondrial superoxide and cytosolic ROS, as well as the largest decrease in ATP (Figure 3B, 3C). NDUFA8 knockdown may result in decreased levels of other complex I subunits responsible for electron transfer(35), or may eliminate important high-turnover sites of oxidation within complex I(9), either of which could contribute to elevated mitochondrial superoxide. Other complex I subunits disproportionately impacted either ATP, mitochondrial superoxide, or cytosolic ROS. Complex I subunits have been annotated by their presence in functional “modules”, based on their roles in either the oxidation of NADH to NAD+ (N-module), reduction of ubiquinone (Q-module), or proximal or distal proton pumping across the mitochondrial membrane (PP- or PD-modules, respectively)(36). However, comparison of the effects of subunits from these different modules on ATP or ROS phenotypes revealed few distinctions except a significant difference between the PP and Q-modules’ cytosolic ROS phenotypes (Figure 3B, Figure S2A). A role for the PP module in regulating ROS is unexpected because the PP module doesn’t contain subunits with iron-sulfur clusters, and also doesn’t directly participate in electron shuttling. It is possible that some PP module subunits like NDUFA8 may impact sites of ROS generation in complex III, which can contribute to cytosolic pools(37, 38), or negatively impact antioxidant systems. Fourteen complex I subunits are “core” subunits that are highly conserved from bacteria to humans and considered to be most critical to function(39-41). There were no significant differences in ATP or ROS phenotypes between core and supernumerary accessory subunits, which generally have less-defined roles in complex I function (Figure S2B), suggesting that more nuanced subunit-specific effects determine how complex I regulates ATP and ROS.

**Fig. 3.**
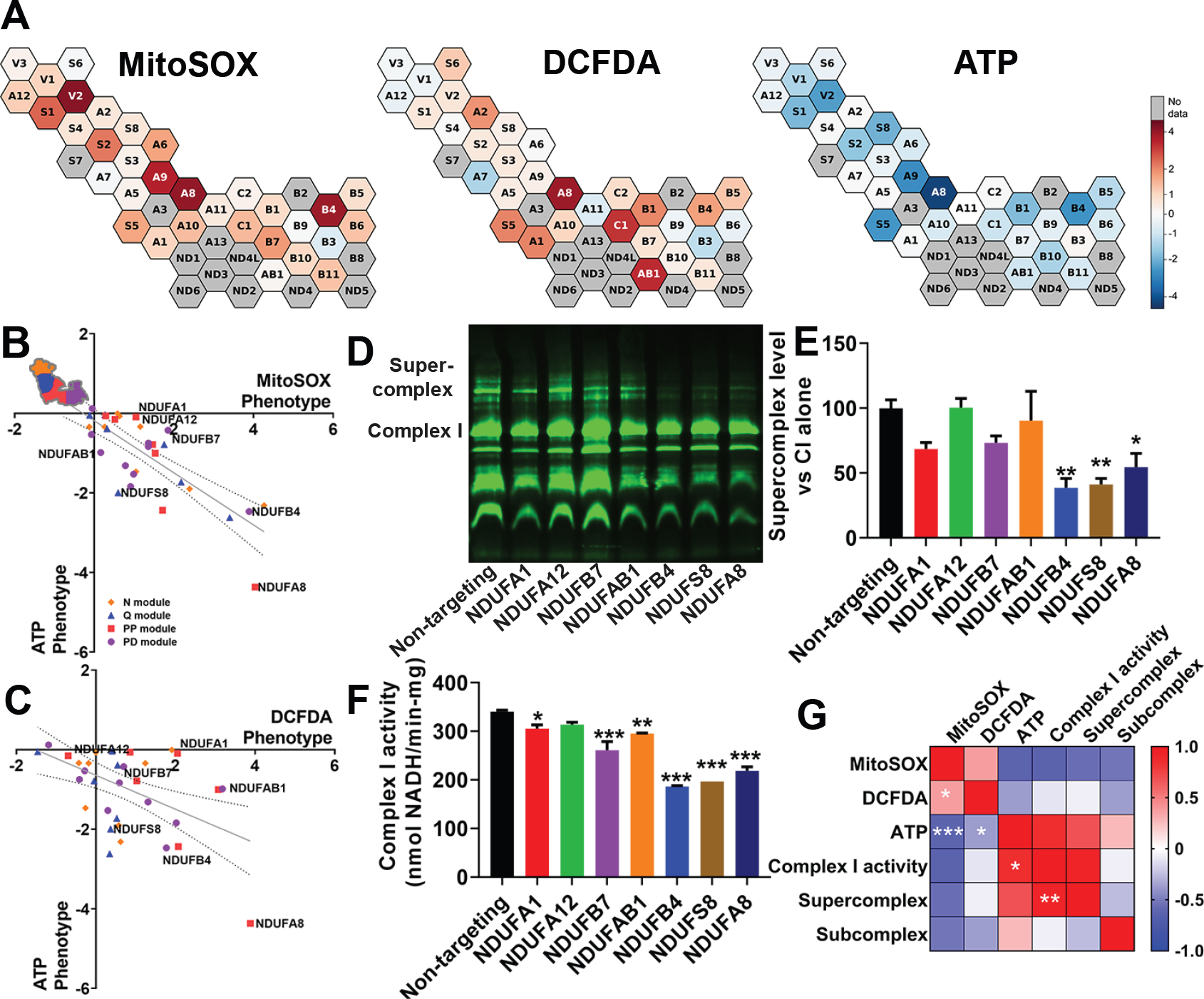
Supercomplex and ATP levels are associated with complex I enzyme activity. A) Cells with complex I subunit knockdown, part of an expanded CRISPRi knockdown library, exhibit a range of mitochondrial ROS or cytosolic ROS phenotypes, as well as a range of ATP phenotypes (previously measured and published in (Bennett et al.(13)), indicated by the color bar. B,C) Complex I subunits grouped by module. The trendline shows linear regression across complex I subunits, and the dotted lines 95% confidence intervals. Subunits are colored based on their association with functional modules within complex I. The notated complex I subunits have prominent effects on ATP and ROS and are examined in follow-up studies. Data are means ± SEM. D) A representative blue-native PAGE gel, loaded with isolated mitochondria for each of seven cell lines expressing CRISPRi knockdown of complex I subunits and a non-targeting control and stained with total OXPHOS human western blot antibody cocktail. E) Quantification of n = 3 blue-native PAGE gels shows that knockdown of NDUFB4, NDUFS8, and NDUFA8 decreases supercomplex levels. F) Quantification of complex I enzyme activity from isolated mitochondria reveals significantly decreased enzyme activity with NDUFA1, NDUFB7, NDUFAB1, NDUFB4, NDUFS8, and NDUFA8 knockdown, from n = 3 replicates. G) Pearson r correlation matrix of mitochondrial ROS, cytosolic ROS, and ATP phenotypes compiled from 33 complex I subunit knockdowns included in CRISPRi screens from n = 2 replicates, along with complex I activity, supercomplex level, and subcomplex level compiled from seven complex I subunit knockdowns from n = 3 replicates. ATP levels were negatively correlated with mitochondrial ROS and cytosolic ROS, and positively correlated with complex I activity. Complex I activity was positively correlated with supercomplex levels. Mitochondrial ROS and cytosolic ROS were also positively correlated. *p<0.05, **p<0.01, ***p<0.001 by one-way ANOVA with Dunnett’s multiple comparisons test (E, F) and Pearson correlation test (G)

To explore these roles, we generated individual cell lines with sgRNA knockdowns targeting seven complex I subunits (NDUFA1, NDUFA12, NDUFB7, NDUFAB1, NDUFB4, NDUFS8, NDUFA8; highlighted in Figure 3B and 3C) (Figure S2C). With the exception of NDUFS8, these subunits are all considered “accessory” or supernumerary subunits. We validated that NDUFA8 knockdown increased mitochondrial ROS, measured using mitochondrial matrix-targeted HyPer7(42) (Figure S2D). We isolated mitochondria from these cell lines and quantified supercomplex levels, subcomplex levels (complexes smaller than the size of fully assembled complex I), and complex I activity. Knockdown of three subunits in particular—NDUFA8, NDUFB4, and NDUFS8—significantly decreased levels of complex I-containing supercomplex relative to assembled complex I levels (Figure 3D, Figure 3E, Figure S2E). Knockdown of these genes, in addition to NDUFA1, NDUFB7, and NDUFAB1, also significantly decreased complex I enzyme activity (Figure 3F). None of the knockdowns significantly increased levels of partially assembled complex I relative to assembled complex I (Figure S2F, Figure S2G), suggesting that the decrease in complex I present in supercomplex is not driven by defects in the assembly process. Notably, knockdown of NDUFA8, NDUFB4, and NDUFS8 had the lowest ATP phenotypes of the subunits examined (Figure 3A, B). Therefore, we investigated if there was a correlation between ATP phenotypes and supercomplex levels upon loss of specific complex I subunits. Indeed, ATP phenotype was highly and significantly correlated with enzyme activity, which in turn was highly and significantly correlated with complex I-containing supercomplex level (Figure 3G, Figure S2H). In contrast, both mitochondrial superoxide and cytosolic ROS levels only weakly correlated with ATP, enzyme activity, or supercomplex level. This analysis also revealed that some complex I subunits have a significant, disproportionately large impact on ATP relative to their impact on supercomplex level or enzyme activity. Notably, NDUFA1 knockdown maintained higher enzyme activity (Figure S2F) and ATP levels (Figure S2I) than would be expected based on its effects on supercomplex levels. This may indicate that the composition of the remaining supercomplexes upon NDUFA1 knockdown are particularly effective in preserving complex I activity and ATP levels. Although cells with NDUFA1-deficient supercomplexes maintained a relatively higher than expected ATP level, this still resulted in increased ROS levels (DCFDA phenotype z-score: 2.81, MitoSOX phenotype z-score: 1.80, ATP phenotype z-score: -0.33). In contrast to NDUFA1, loss of NDUFA8 decreased ATP to a greater extent than expected from its impact on supercomplex levels (Figure S2I). These results demonstrate a systems-level trend across complex I subunits where supercomplex levels and enzyme activity correlate strongly with ATP level, and where ROS levels are regulated uniquely across different complex I subunits. Our findings also identify specific complex I subunits whose expression are critical to maintaining cellular energy, and may be therapeutic targets for preservation in diseases of energy failure.

### Identification of ATP-independent regulators of ROS

In order to identify genetic regulators of ROS that were independent of overall ATP levels, we examined knockdowns that resulted in high- and low-mitochondrial superoxide but did not alter ATP phenotypes. Interestingly, genes related to peroxisomal ether lipid synthesis were associated with both high- and low-ROS phenotypes (Figure 2C, Figure 4A). Ether lipids, which include plasmalogens, are glycerophospholipids synthesized in peroxisomes known to act as endogenous antioxidants and to regulate membrane structure and fluidity and signal-transduction(43). Knockdown of peroxisomal ether lipid synthesis genes (PEX5, PEX7, AGPS, FAR1, GNPAT, MitoSOX phenotype z-scores: -4.32, -4.58, -3.39, -3.33, -2.20, respectively) decreased mitochondrial ROS, while knockdown of the mitochondrial enzyme dihydroorotate dehydrogenase (DHODH) increased mitochondrial ROS. Inhibition of DHODH has been shown to increase supercomplex levels by increasing peroxisomal ether lipid synthesis(44). To test the dependence of ATP, ROS and supercomplex levels on DHODH inhibition and peroxisomal ether lipid synthesis, we first cultured cells in respiration-only media in the presence of vehicle or DHODH-inhibitors (brequinar or vidofludimus) under conditions shown to influence ether lipid synthesis and supercomplex formation(44). Interestingly, DHODH inhibition did increase mitochondrial superoxide measured by either MitoSOX or MitoNeoD (Figure 4B, Figure 4C, Figure 4D, Figure 4E), but did not consistently increase ATP level (Figure 4F, Figure 4G), consistent with a lack of effect in our ATPome analysis(13). Next, we tested the effect of ether lipid precursor sn-1-O-hexadecylglycerol (OHG), which can increase ether lipid levels(45), and found that it increased mitochondrial superoxide in cells expressing a control, non-targeting CRISPRi sgRNA (Figure 4H, Figure 4I). However, OHG did not increase mitochondrial superoxide in cells with NDUFS8 knock down (Figure 4H, Figure 4I). These data indicate that complex I subunits that are critical for supercomplex assembly are also required for ether lipid modulation of mitochondrial ROS.

**Fig. 4.**
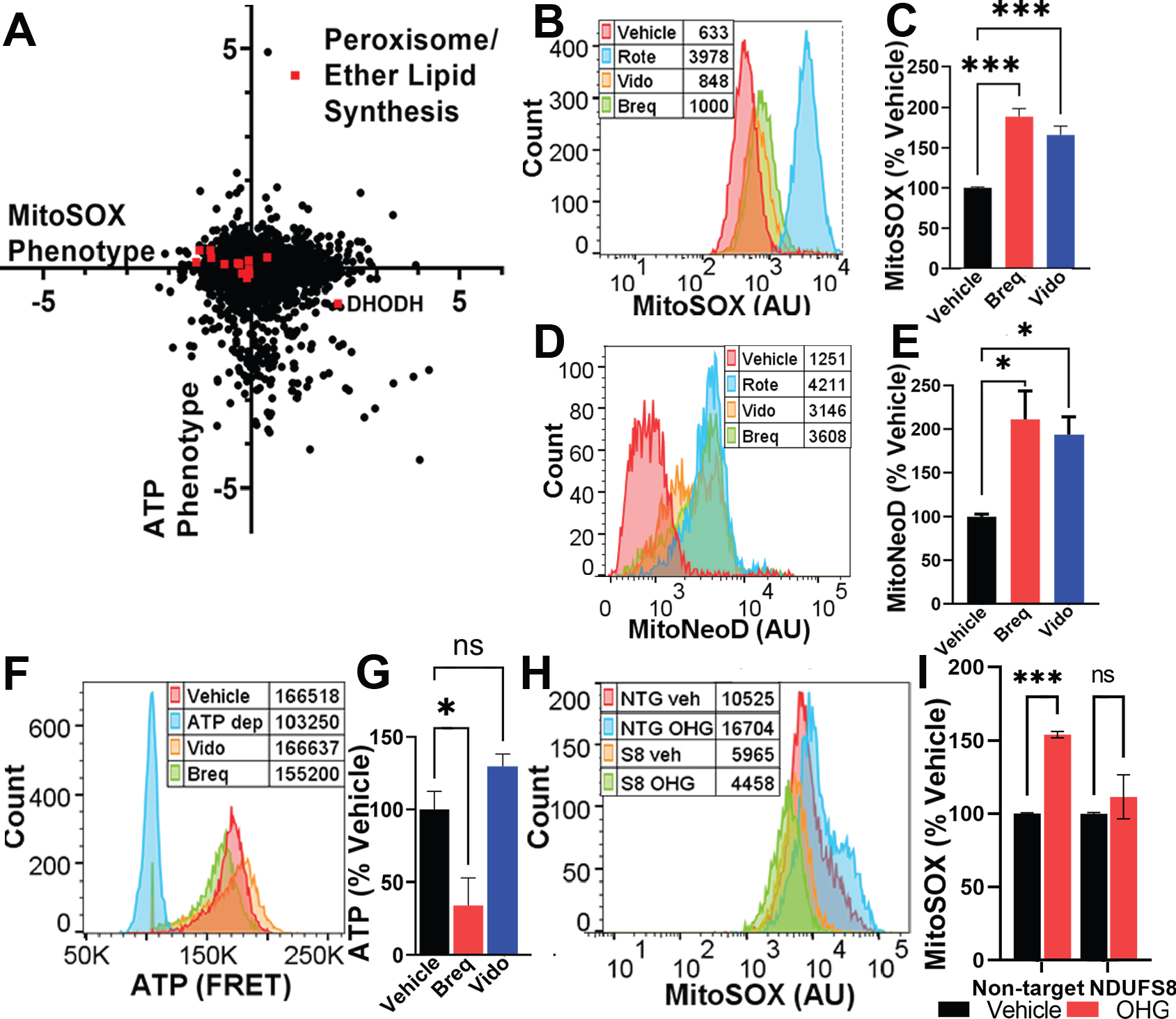
Ether lipids bi-directionally control mitochondrial ROS. A) Peroxisomal and ether lipid synthesis genes and DHODH, whose inhibition elevates peroxisomal ether lipid levels(44), span a wide range of mitochondrial ROS phenotypes, without impacting ATP. Data compiled from n = 2 replicates. B,C) Representative plot with peak averages and quantification of cells incubated with DHODH inhibitors vidofludimus (Vido, 10 µM), brequinar (Breq, 0.5 µM), or rotenone (Rote, 2 nM) exhibit elevated mitochondrial ROS, measured with MitoSOX. n = 2 replicates with ≥ 10,000 cells per condition. D,E) Similar increases in mitochondrial ROS were observed with DHODH inhibitors, measured with MitoNeoD. n = 2 replicates with ≥ 10,000 cells per condition. F,G) Representative plot with peak averages and quantification of cells expressing a FRET-based sensor for ATP measured using FACS. There was no consistent difference in ATP in response to 48 h incubation with DHODH inhibitor drugs. ATP-depleted (ATP dep) control cells were treated with oligomycin (10 µM) and 2DG(10 mM). n = 2 replicates with ≥10,000 cells per condition. H,I) Cells expressing a non-targeting CRISPRi sgRNA (NTG) exhibit elevated mitochondrial ROS with DHODH at 24 h incubation with 20 µM ether lipid precursor sn-1-O-hexadecylglycerol (OHG). In contrast, K562 cells with CRISPRi knockdown of NDUFS8 (S8) have a diminished mitochondrial ROS response to ether lipid precursor. n = 2 experiments with ≥10,000 cells per condition. ns = not significant, *p < 0.05, ***p < 0.001 by one-way ANOVA.

## Discussion

Here, we used a CRISPR- and FACS-based screening platform to identify genes and pathways that regulate ROS levels. We combined these results with those of prior parallel screens against ATP to delineate ATP- and ROS-regulating hits, and identified genes and metabolic conditions that decouple energy production from oxidative stress. The analyses reveal critical subunits for respiratory chain function, identify a novel regulator of mitochondrial superoxide, and elucidate a systems-wide connection between complex I activity, supercomplex and ATP levels.

Several primary sites for mitochondrial as well as non-mitochondrial ROS production have been identified(46-48). Our studies align with previous reports of the major sites of mitochondrial ROS generation, particularly complexes I and III in the mitochondrial electron transport chain(1). Mutations in these complexes and in CoQ10 biosynthesis are known to cause inborn mitochondrial disorders like Leigh syndrome(49), and we observe that knockdown of genes associated with these complexes and pathways most frequently increased mitochondrial ROS and decreased ATP. Notably, knockdown of specific complex I genes in the respiratory chain elevated mitochondrial and cytosolic ROS. In the latter case, mitochondrial superoxide is presumably converted into H_2_O_2_ that diffuses out of the mitochondria to increase cytosolic ROS levels. It is unclear why knockdown of certain complex I subunits increased both mitochondrial and cytosolic ROS, while knockdown of other complex I subunits increased only mitochondrial ROS. However, given the enrichment of stronger mitochondrial ROS hits among those that also increased cytosolic ROS, it may simply reflect cells with greater levels of mitochondrial ROS which will have a higher efflux into the cytosol.

We found that the cholesterol biosynthesis pathway also strongly impacted both mitochondrial superoxide and cytosolic ROS pools. Statins are drugs that inhibit cholesterol biosynthesis, and are widely used to reduce morbidity and mortality from cardiovascular disease. Much work has been devoted to studying the antioxidant and ROS-lowering effects of statins(50), however, our data suggest that inhibiting cholesterol synthesis at the cellular level can elevate mitochondrial superoxide and cytosolic ROS. This is consistent with observations by Bouitbir et al., who showed that skeletal muscle from patients with statin-induced muscular myopathy had elevated ROS production and decreased antioxidant defense(51, 52). It is possible that the ROS-inducing effects of cholesterol biosynthesis inhibition may be caused by other important functions of cholesterol, including the regulation of membrane fluidity(53), which is known to regulate mitochondrial respiratory chain function(54) and increase mitochondrial ROS production and toxicity(55).

Despite established effects of ROS in damaging cellular components, antioxidant therapies have been largely ineffective in clinical studies(56). In recent studies, we showed that ATP deficits resulting from COX11 knockdown, and in cells derived from patients with a COX11 mutation, were restored by CoQ10 supplementation(28) (57). The current study builds on this approach, offering, to our knowledge, the first systems-level analysis of antioxidant efficacy. Our findings indicate that antioxidant efficacy may depend on the context of ROS dysregulation. For example, we found that cells with increased ROS due to knockdown of CoQ10 biosynthetic genes were particularly responsive to different antioxidant therapies.

Complex IV is not considered a direct source of ROS production. However, we demonstrate on a systems level that knockdown of complex IV assembly factors (COX11, SCO1, COX16 and COX18) decreased ROS levels, in addition to decreasing ATP(13, 28). The mechanism for this effect is unknown, although it is notable that complex IV is required for complex I stability or assembly, and hence complex I levels may be markedly decreased in the absence of complex IV(58-60), perhaps leading to less superoxide production from complex I and distal sites on the respiratory chain. Similar to the effects on HK2 knockdown cells discussed above, we observed that MitoQ treatment also increased mitochondrial ROS in COX11 knockdown cells (Dataset S1). Guarás et al. observed that saturation of the CoQ10 pool with electrons results in destabilization of complex I (61). Since MitoQ treatment likely increases mitochondrial CoQ10 pool size and oxidation capacity, the increased mitochondrial superoxide observed with MitoQ treatment of COX11-deficient cells may be dependent on increased stability and activity of complex I. Importantly, our data suggest that for patients with COX11 mutation(57), elevated ROS would not be expected, and loss of ATP may be a more important driver of disease. Similarly, we found that knockdown of mitochondrial ribosomal proteins predominantly impacts ATP and not ROS. Therefore, based on our previously published work(28), a promising therapeutic approach for individuals with mutations in these genes may be changing available metabolic substrate or diet to favor glycolytic energy production.

Mutations in complex I are the most common inherited defects in the respiratory chain associated with mitochondrial diseases (32-34), however, the functional effects of loss of individual subunits is not well understood. We observed that knockdown of complex I subunits spanned a wide range of ATP and ROS phenotypes that were generally not explained by location in functional “modules”(36). In order to better understand if and how loss of individual complex I subunits can decouple ATP and ROS production, we measured complex I activity and supercomplex levels in a range of complex I knockdown lines. Our data reveal a high correlation between the degree of complex I subunit knockdown and deficits in enzyme activity, supercomplex levels, and ATP level. Most notably, knockdown of the three subunits that resulted in the lowest ATP levels—NDUFA8, NDUFB4, and NDUFS8—also resulted in the lowest supercomplex levels and complex I enzyme activity. Complex I has been observed to adopt two catalytically and structurally different states, an active and de—active conformation(4). The mechanism for the active-de-active transition is known to involve conformational rearrangements of NDUFA9 and ND1, and the oxidation state of ND3(62, 63), all located in the vicinity of the quinone binding site(5). We speculate that loss of some of the complex I subunits that were associated with low ATP, and particularly with low enzyme activity like NDUFA8, may push complex I towards the de-active conformation.

The assembly of respiratory chain complexes into supercomplexes or respirasomes has been hypothesized to offer structural or functional advantages, possibly including enhanced stability or resistance to degradation, improved electron transport efficiency and substrate channeling, and decreased leak of electrons to produce ROS(64). In support of this model, Lopez-Fabuel et al. observed that knockdown of NDUFS1 increased ROS and decreased supercomplex level, while overexpression of NDUFS1 increased supercomplex formation and decreased ROS(10). Indeed, our data show that NDUFS1 knockdown is associated with increased ROS as well as decreased ATP (MitoSOX phenotype z-score 4.32, ATP phenotype z-score -9.29), and based on our systems-level analysis, would be expected to decrease supercomplex formation. Thus, maintaining supercomplex levels and enzyme activity by preserving expression of multiple specific complex I subunits may be a strategy to address energy deficits.

We identified ether lipids, a class of glycerophospholipids with an alkyl chain attached by an ether bond at the *sn-* 1 position, as a robust way to bi-directionally modulate mitochondrial ROS. We found that knockdown of ether lipid biosynthetic genes resulted in lower levels of mitochondrial superoxide, while knockdown of the mitochondrial enzyme DHODH, which increases supercomplex levels(44) by inducing peroxisomal production of ether lipids, increased mitochondrial superoxide. In our system, we observed that both DHODH inhibitors and ether lipid precursor increase mitochondrial superoxide. However, this increase could be blocked by NDUFS8 knockdown, suggesting that this subunit, or perhaps intact functional complex, is required for the ether lipid effects on supercomplex formation and subsequent mitochondrial ROS production. Ether lipids are believed to regulate membrane fluidity(43) and curvature(44), which may create stable regions for respiratory chain complex association into supercomplexes, but could also alter points of electron transfer and increase levels of ROS production. Ether lipid synthesis inhibitors are being explored as potential cancer therapeutics(65), which may point to the benefit of lowering mitochondrial ROS in the treatment of cancer. Interestingly, Jain et al. observed that knockdown of ether lipid biosynthetic genes impaired cell survival in hypoxic conditions(54), raising the possibility that ether lipid-mediated assembly of supercomplexes, and supercomplex-derived mitochondrial superoxide, are important parts of cellular adaptation to hypoxic conditions.

In summary, our findings expand our understanding of the function of the respiratory chain and supercomplexes, reveal distinct systems-level regulation of ATP and ROS levels, and suggest that antioxidant therapies should be carefully chosen based on the site of ROS biogenesis. Importantly, we demonstrate that loss of complex I subunits can decrease complex I activity, supercomplex levels, and ATP levels, without necessarily increasing ROS. We furthermore identify ether lipids as a dynamic regulator of mitochondrial ROS, which fits with previous observations connecting ether lipids and supercomplex formation. Future studies are required to determine if ether lipids promote the formation of specific stoichiometries of supercomplex, and to elucidate the causal connections between supercomplex level, complex I activity, and ATP level. Preservation of specific supercomplex stoichiometries, either through manipulation of mitochondrial lipids or stabilizing individual subunits, may be a promising avenue to tune cellular ATP and ROS.

## Materials and Methods

### Cell Lines

These studies were conducted with K562 human female leukemia cells, which were provided by the lab of Jonathan Weissman, similar to those used in our previously published ATP screen(13). The cells had stable integration of dCas9-KRAB enabling CRISPRi. For cell culture, K562 cells were maintained in RPMI-1640 media with 25mM HEPES, 0.3g/L L-glutamine, 2.0 g/L NaHCO_3_, and supplemented with 10% fetal bovine serum, 100 units/mL penicillin, 100 mg/mL streptomycin, and 2 mM glutamine.

### ROS FACS screen and analysis

A detailed protocol for FACS screening to detect ROS can be found at https://doi.org/10.17504/protocols.io.e6nvwdz87lmk/v1. We created a mini-library of individual sgRNAs targeting ATP-impacting hits based on our previously published work, in order to more rapidly conduct analysis with fewer cells. This mini-library was comprised of 1-3 sgRNAs per gene based on robust ATP phenotype. This mini-library included a total of 450 sgRNAs, 19 non-targeting sgRNAs as controls and 431 sgRNAs targeting 219 genes. First, two complementary oligonucleotide sequences per sgRNA were annealed and individually ligated into a lentiviral backbone plasmid used by Gilbert et al(66). Full CRISPRi sgRNA sublibraries were provided by the lab of Jonathan Weissman. After confirming sequences of individual clones with Sanger sequencing, lentivirus was created by the UCSF Viracore and transduced into dCas9-KRAB expressing K562 cells with polybrene (8 µg/mL) via spinfection. After two days, cells were maintained in puromycin (0.65 µg/mL) for 5 days to select for cells expressing sgRNA.

Cells were resuspended in PBS with metabolic substrate and ROS dyes (2.5 µM MitoSOX or 5 µM MitoNeoD) for 30 min prior to flow cytometry. Most studies were conducted in respiratory conditions (2% fetal bovine serum, 10 mM pyruvate, 10 mM 2-deoxyglucose) in order to identify drivers of mitochondrial ROS when cells are dependent on respiration for ATP. Where indicated, cells were resuspended in glycolytic conditions (2% fetal bovine serum, 2 mM glucose, 5 µM oligomycin, 3 mM 2-deoxyglucose) or basal conditions (2% fetal bovine serum, 10 mM glucose, 5 mM pyruvate). In studies with antioxidants, cells were incubated with either 10 µM MitoQ, or 1 mM Trolox for 2 hours prior to cell sorting. All flow cytometry and cell sorting was conducted on either a BD FACSAria II (RRID:SCR_018934) or a BD FACSAria Fusion (facility RRID:SCR_021714).

Cells sorted by FACS were centrifuged and kept at -20 °C until further processing, which has been described previously (13, 28), but is summarized as follows. Genomic DNA was isolated from cell pellets using the Macherey-Nagel NucleoBond Xtra Midi Plus kit. sgRNAs incorporated via lentivirus into genomic DNA were amplified and affixed with sequencing adapters and barcodes in a single PCR step, with 1.5 µg undigested genomic DNA per PCR reaction using Q5 HotStart High Fidelity Polymerase (New England Biolabs). Resulting PCR product from multiple reactions per sample were pooled. PCR product was separated from unincorporated primers using the GeneRead Size Selection Kit. Purity and quality of the PCR product was measured using a bioanalyzer (Agilent, RRID:SCR_018043). Finally, sequencing was performed on an Illumina HiSeq 2500 (RRID:SCR_016383).

### Confirmation of gene knockdown

A detailed protocol for confirming gene knockdown with RT-PCR can be found at https://doi.org/10.17504/protocols.io.dm6gp3yo1vzp/v1. Relative gene expression quantifications were performed using qRT-PCR on a 7900HT Fast Real-Time PCR System (Applied Biosystem; RRID:SCR_018060), using VIC-MGB human ACTB as an endogenous control (β-actin, ThermoFisher #4326315E) and FAM-MGB TaqMan Gene Expression Assays (ThermoFisher, assay ID: Hs00204417_m1-NDUFA8, Hs00244980_m1-NDUFA1, Hs00984333_m1-NDUFA12, Hs00958815_g1-NDUFB7, Hs00192290_m1-NDUFAB1, Hs00853558_g1-NDUFB4, Hs00159597_m1-NDUFS8, Hs02578754_g1-NDUFS5). cDNA and PCR reactions were prepared using the Cells-to-CT kit (ThermoFisher), following their protocol with a standard reverse transcription cycle (37 °C for 6 minutes, followed by 95 °C for 5 minutes and holding at 4 °C), and standard qRT-PCR conditions (UDG incubation of 50 °C for 2 minutes, enzyme activation at 95 °C for 10 minutes, and PCR cycles of 95 °C for 15 seconds, 60 °C for 1 minute, repeated for 40 cycles). All reactions were performed in a 384-well plate format in duplicate and compiled from at least 2 independent experiments. Fold changes in expression were calculated using the 2^-ΔΔCT^ method.

### Functional effects of individual gene knockdown

A detailed protocol for characterizing isolated mitochondria can be found at https://doi.org/10.17504/protocols.io.e6nvwdz87lmk/v1. To investigate the functional impacts of individual sgRNA knockdowns on mitochondrial function, we first digested mitochondrial matrix-localizing HyPer7 (pCS2+MLS-HyPer7, RRID:Addgene_136470) with BamH1 and XhoI and ligated into our lentiviral backbone (13), before transduction into CRISPRi capable K562 cells. Cells expressing the sensor were enriched by flow cytometry, before transducing with non-targeting or NDUFA8. We also created individual K562 cell lines expressing other individual complex I-targeting sgRNAs. After culturing 100 million cells, crude mitochondria were isolated by using a Wheaton Dounce tissue homogenizer. Mass of isolated mitochondria were measured using a Pierce BCA protein assay kit.

We measured enzyme activity of complex I by dispensing 10 µg of crude mitochondria preps into a 96-well assay plate containing 50 mM pH 7.4 potassium phosphate buffer, 1 mM EDTA, 2 mM potassium cyanide, 2 mM sodium azide, 2.5 mg/mL bovine serum albumin, 5 mM MgCl_2_, 2 µM Antimycin A, 100 µM decylubiquinone, and 300 µM K_2_NADH, with or without 1 µM rotenone. Immediately after combining mitochondria with the above reagents, 340 nm absorbance was measured for a period of 45 minutes on a Spectramax M4 platereader.

To quantify levels of supercomplex (67), we solubilized 50 µg of crude mitochondria preps with 5% digitonin at a 8g/g digitonin to protein ratio. Solubilized mitochondria were loaded onto a Native PAGE 3-12% gradient gel, before transferring to PVDF membranes with an iBlot Gel Transfer Device. PVDF membranes were blocked and stained with total OXPHOS human western blot antibody cocktail (Abcam Cat# ab110411, RRID:AB_2756818, 1:500). Blots were visualized using an Odyssey infrared imaging system (RRID:SCR_022510) and then analyzed with Image Studio Lite Software (Li-cor Biosciences; RRID:SCR_013715).

### Quantification and statistical analysis

An individual sgRNA’s ROS phenotype was calculated as the fold enrichment in the high versus low ROS fractions. For the smaller sgRNA library, correlation analysis for targeting and non-targeting guides across replicates or with antioxidant treatment were quantified with linear regression and comparison of slopes to a hypothetical value of 0 or 1 via extra sum-of-squares F-test. ROS phenotypes of individual classes of sgRNA within the smaller library were compared to non-targeting sgRNA using one-way ANOVA with Dunnett’s multiple comparisons test. For the larger scale sgRNA library, a given gene’s perturbation phenotype was calculated as the average enrichment of the strongest of three sgRNA guides. For each gene, we also calculated a Mann-Whitney p-value, where a lower p-value corresponded to greater agreement in phenotype between the ten sgRNAs targeting each gene.

Pathway-wide enrichment among the screened genes was measured by first assembling lists of genes ranked by their ROS phenotypes. These lists were uploaded to the Galaxy web platform (RRID:SCR_006281), and the public server at usegalaxy.org and fast pre-ranked fast gene set enrichment analysis (68) were used to analyze the data (69). Default settings, including a minimum and maximum gene set size of 1 and 500, respectively, and one thousand random sample permutation using the Molecular Signatures Database c2 v7.4 and c5 v7.4 (RRID:SCR_016863).

Comparisons between individual cell lines’ supercomplex level, enzyme activity, and subcomplex level, or between core and supernumerary or complex I modules were performed using one-way ANOVA with Dunnett’s multiple comparisons test. Correlation matrices were calculated with Pearson r scores, with p values calculated using t-tests.

### Data, Materials and Software Availability

All data produced for this study are included in the article, or included in supplementary information. Datasets are available at: https://doi.org/10.5281/zenodo.8419489.

## ACKNOWLEDGEMENTS

We thank Gwyneth Hutchinson for technical assistance and Isha Jain for critical reading of manuscript. We thank Brandon DeSousa for his help plotting figures. We thank Eric Chow and the UCSF Center for Advanced Technology, as well as Kathryn Claiborn for helping edit the manuscript, Giovanni Maki for help with graphics and Erica Delin for administrative assistance. This work was funded in part by R01 AG065428 (K.N.), the UCSF Bakar Aging Research Institute (BARI, K.N.), and by Aligning Science Across Parkinson’s [ASAP-020529] through the Michael J. Fox Foundation for Parkinson’s Research (MJFF). It was also supported by a Berkelhammer Award for Excellence in Neuroscience (N.B.), F32 AG063457-02 (N.B.), K01AG078485 (N.B.), and R01 AGO68091 (A.L.O.). We thank the Gladstone Flow Cytometry Core for assistance and use of flow cytometer equipment, supported by NIH S10RR028962, NIH P30 AI027763, and the James B. Pendleton Charitable Trust. For the purpose of open access, the author has applied a CC BY public copyright license to all Author Accepted Manuscripts arising from this submission.

## Figure Legends

**Fig. S1**. Antioxidant effect on ROS levels depends on deficient pathway. A) ROS phenotypes of K562 cells expressing a mini-library of CRISPRi sgRNA detected with MitoNeoD were similar to those measured with MitoSOX, with some of the same hit genes like NDUFA8, and similar patterns with complex IV-associated gene knockdowns having low ROS. B) Some of these same patterns were observed in cells treated with 0.1 µM Decyl-TPP, the control for MitoQ Treatment. Cells treated with either C,D) low dose of MitoQ (0.1 µM) or E) high dose (10 µM) of MitoQ respond differently depending on the genes knocked down. MitoQ at both doses abrogated the increase in ROS from CRISPRi knockdown of CoQ10 biosynthetic genes. In contrast, 10 µM MitoQ treatment increased mitochondrial and cytosolic ROS following knockdown of HK2 and VDAC1. F) Trolox (1 mM) abrogated the increase in ROS following knockdown of CoQ10 biosynthetic genes. Trolox treatment also decreased ROS levels across all targeting CRISPRi sgRNA in aggregate. n = 2 replicates. G) Pearson r correlation matrix of mitochondrial content, ATP level and ATP depletion phenotypes in basal, respiration-only, and glycolysis-only conditions collected previously from n = 2 replicates(13), along with ROS phenotypes in the same conditions in cells expressing CRISPRi knockdown libraries. Across ATP level phenotypes, there was only significant correlation between basal and glycolysis-only conditions. In contrast, ROS levels correlated between all substrate conditions. *p < 0.05, **p < 0.01, ***p < 0.001 by one-way ANOVA with Dunnett’s multiple comparisons test (A,B) and (C) by Pearson correlation test.

**Fig. S2**. CRISPRi knockdown of complex I subunits decreases supercomplex levels. A) Knockdown of subunits within functional modules of complex I did not significantly affect either mitochondrial ROS or ATP. Knockdown of subunits within the Q module and PP module significantly differed in their effects on cytosolic ROS. B) Effect of knockdown of core conserved subunits did not significantly differ from the effect of knockdown of non-core or supernumerary subunits on mitochondrial ROS, cytosolic ROS, or ATP. Data compiled from n = 2 experiments. ATP phenotype data previously collected in Bennett et al (13). C) RT-qPCR of K562 lines expressing CRISPRi sgRNA knocking down individual complex I subunits. n = 2-10 replicates per cell line. D) Knockdown of NDUFA8 in K562 cells increases mitochondrial ROS, as measured by mitochondrial matrix-targeted HyPer7. Data compiled from n=2 experiments. E) Additional representative blue-native PAGE gel replicate, loaded with isolated mitochondria for each of seven cell lines expressing CRISPRi knockdown of complex I subunits and a non-targeting control, and stained with total OXPHOS human western blot antibody cocktail. F) Representative blue-native PAGE gel, loaded with isolated mitochondria for each of seven cell lines expressing CRISPRi knockdown of complex I subunits and a non-targeting control, and stained with NDUFS4-targeting antibody to identify NDUFS4-containing sub-complexes at molecular weights less than fully assembled complex I. G) There were no significant differences in subcomplex levels versus non-targeting controls. n = 2 blue-native PAGE gels Supercomplex levels correlate with complex I activity (H) and ATP levels (I). NDUFA1 knockdown has proportionally higher complex I activity and ATP than other subunits, given its level of supercomplex, indicating that its knockdown may preserve a supercomplex species of higher energetic productivity. Trendline shows linear regression across the complex I knockdown cell lines, and the dotted lines are 95% confidence intervals. ATP phenotype data previously collected in Bennett et al (13). ns = not significant, *p < 0.05, **p < 0.01, ***p < 0.001 by 2-way ANOVA with Tukey’s (A,B), or Bonferroni’s (C) multiple comparisons test, Student’s t-test (D), and one-way ANOVA with Dunnett’s multiple comparisons test (G).

**Table S1**. ROS phenotypes are robust and sensitive to antioxidants. The first column indicates which plot in Figure 1 corresponds to the listed statistics. For both ROS dyes, we find that the slopes are significantly non-zero, indicating significant correlation and robustness, by extra sum-of-squares F-test between phenotypes, n=2 replicates. We also find that for all antioxidant treatments on all ROS dyes, there is a significant effect of the antioxidants on lowering ROS phenotypes for targeting guides by extra sum-of-squares F-test between vehicle and antioxidant treatment, n=2 replicates. Notably, the MitoQ versus vehicle treatment for targeting guides have slopes significantly different than 0, while Trolox versus vehicle are not, indicating that Trolox had a stronger antioxidant effect across all targeting guides.

**Dataset S1**. Impact of CRISPRi knockdown and antioxidants on ROS phenotypes. K562 cells expressing a mini-library of CRISPRi guides were treated with ROS dyes (MitoSOX, DCFDA, or MitoNeoD) and then sorted by FACS into high- and low-ROS fractions. ROS phenotypes were quantified by measuring guide enrichment in the high versus low ROS fractions (log2 fold-representation scale). Blank cells correspond to guides that did not pass minimum representation thresholds in either cell fraction for a given experimental replicate. N=2 experimental replicates.

**Dataset S2**. ROS phenotypes following genome-scale CRISPRi knockdowns. Mitochondrial and cytosolic ROS were quantified by measuring guide enrichment (log2 fold representation) in the high versus low ROS cell fractions. Mann-Whitney P-values correspond to the degree of agreement between top guides targeting a given gene. Z-scores were calculated relative to the distribution of pseudogenes, derived from non-targeting control guides included in the library of CRISPRi guides. Hits were determined for genes with either z-scores greater than 3 or less than -3, or with Mann-Whitney P-values less than 0.05. N=2 experimental replicates.

