## Supplementary figures and images for "Systems-level analyses dissociate genetic regulators of reactive oxygen species and energy production"

### Figure S1

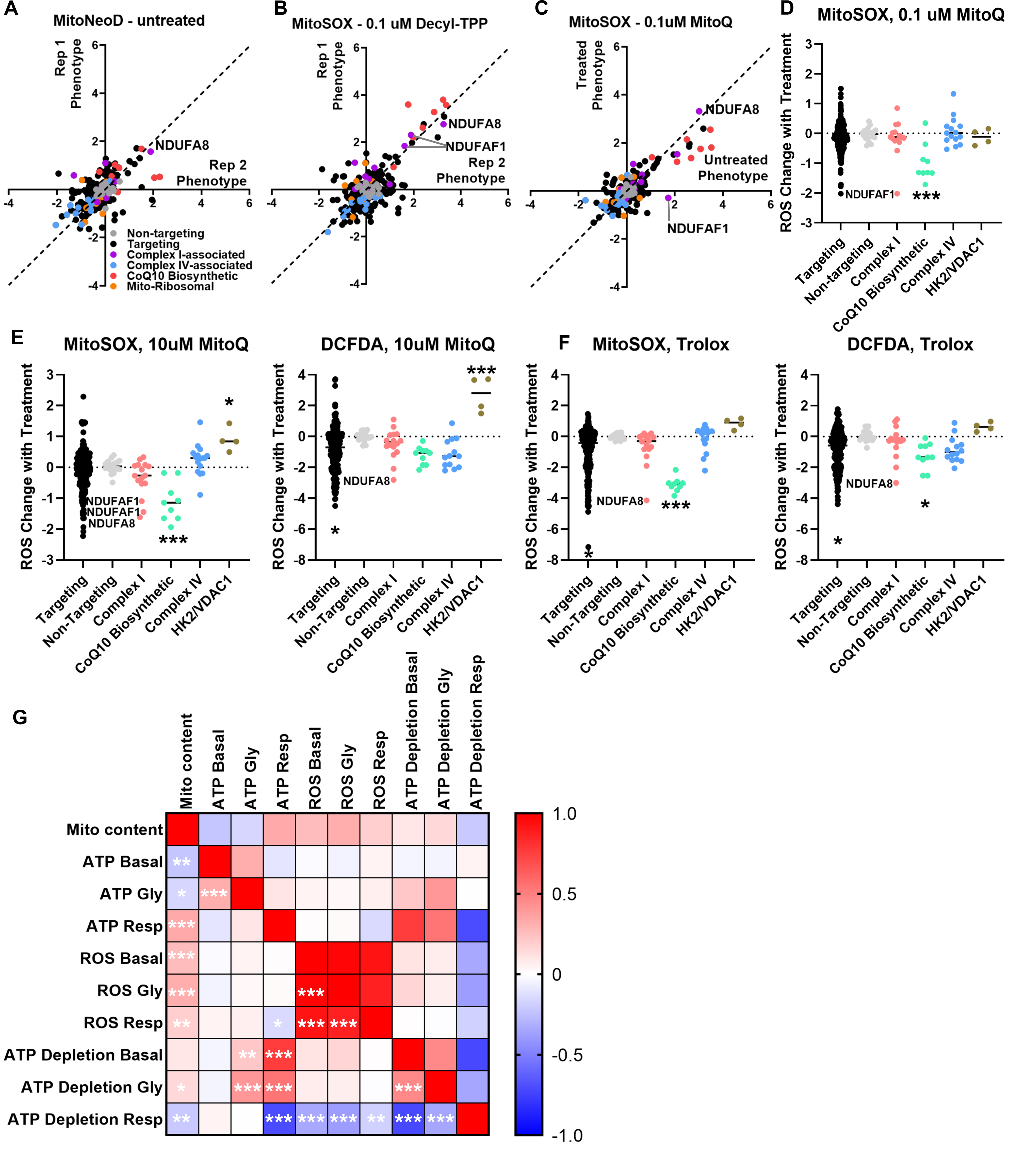

### Figure S2

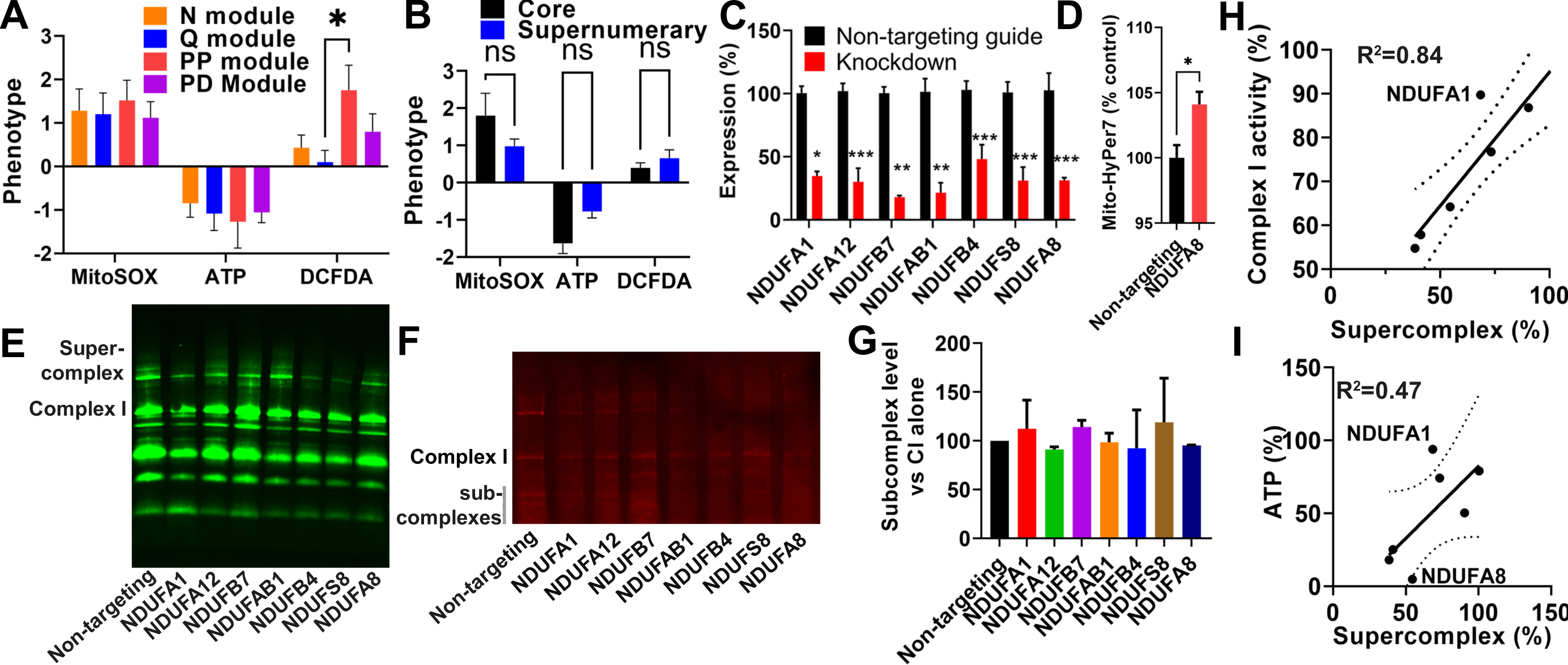
